# Distinct Epigenomic Patterns Are Associated with Haploinsufficiency and Predict Risk Genes of Developmental Disorders

**DOI:** 10.1101/205849

**Authors:** Xinwei Han, Siying Chen, Elise Flynn, Shuang Wu, Dana Wintner, Yufeng Shen

## Abstract

Haploinsufficiency is a major mechanism of genetic risk in developmental disorders. Accurate prediction of haploinsufficient genes is essential for prioritizing and interpreting deleterious variants in genetic studies. Current methods based on mutation intolerance in population data suffer from inadequate power for genes with short transcripts. Here we showed haploinsufficiency is strongly associated with epigenomic patterns, and then developed a new computational method (Episcore) to predict haploinsufficiency from epigenomic data from a broad range of tissue and cell types using machine learning methods. Based on data from recent exome sequencing studies of developmental disorders, Episcore achieved better performance in prioritizing loss of function *de novo* variants than current methods. We further showed that Episcore was less biased with gene size, and was complementary to mutation intolerance metrics for prioritizing loss of function variants. Our approach enables new applications of epigenomic data and facilitates discovery and interpretation of novel risk variants in studies of developmental disorders.

## Introduction

Haploinsufficiency (HIS) due to hemizygous deletions or heterozygous likely-gene-disrupting (LGD) variants plays a central role in the pathogenesis of various diseases. Recent large-scale exome and genome sequencing studies of developmental disorders, including autism, intellectual disability, developmental delay, and congenital heart disease ^1–5^, have estimated that *de novo* LGD mutations explain the cause of a significant portion of patients with these developmental disorders, and the enrichment rate of *de novo* LGD variants indicates about half of these variants are associated with disease risk. However, relatively few genes have multiple LGD variants (“recurrence”) in a cohort ^1,2,6^, lacking of which provides insufficient statistical evidence to distinguish individual risk genes from the ones with random mutations ^7^. On the other hand, most of the enrichment of LGD variants can be explained by HIS genes ^6^. Therefore, a comprehensive catalog of HIS genes can greatly help interpreting and prioritizing mutations in genetic studies.

Currently, there are two main approaches of predicting HIS genes based on high-throughput data. Huang et al. uses a combination of genetic, transcriptional and protein-protein interaction features from various sources to estimate haploinsufficient probabilities for 12,443 genes ^8^. Using similar input information, Steinberg et al. generated the probabilities for more (over 19,700) human genes by a Support Vector Machine (SVM) model ^9^. The other approach is based on mutation intolerance ^10–12^ in populations that do not have early onset developmental disorders. Lek et al 2016 ^11^ estimated each gene’s probability of haploinsufficiency (pLI: Probability of being Loss-of-function Intolerant) based on the depletion of rare LGD variants in over 60,000 exome sequencing samples. Although effective, ExAC pLI is biased towards genes with longer transcripts or higher background mutation rates, since the statistical power of assessing the significance depends on a relatively large expected number of rare LGD variants from background mutations.

We sought to predict HIS using epigenomic data that are orthogonal to genetic variants and generally independent of gene size. Our method is motivated by recent studies indicating that specific epigenomic patterns are associated with genes that are likely haploinsufficient. Specifically, genes with increased breadth of H3K4me3, typically associated with actively transcribing promoters, are enriched with tumor suppressor genes ^13^, which are predominantly haploinsufficient based on somatic mutation patterns ^14^. Another study reported H3K4me3 breadth regulates transcriptional precision ^15^, which is critical for dosage sensitivity. These observations led us to hypothesize that haploinsufficient genes are tightly regulated by a combination of transcription factors and epigenomic modifications to achieve spatiotemporal precision of gene expression, and such regulation can be detected by distinct patterns of epigenomic marks in relevant tissues and cell types. Based on this model, we developed a Random Forest–based method (“Episcore”) using epigenomic data from the Epigenomic Roadmap ^16^ and ENCODE Projects ^17^ as input features and a few hundreds of curated HIS genes as positive training data. To assess the performance of prioritizing candidate risk variants in real-world genetic studies, we used large data sets of *de novo* mutations from recent studies of birth defects and neurodevelopmental disorders and showed that Episcore had better performance than existing methods. Additionally, Episcore is less biased by gene length or background mutation rate and complementary to mutation-based metrics in HIS-based gene prioritization. Our analysis indicates that epigenomic features in stem cells, brain tissues, and fetal tissues contribute more to Episcore than others.

## Results

### Haploinsufficient (HIS) and Haplosufficient (HS) genes show distinct distributions of epigenomic features

To examine the correlation of gene haploinsufficiency and epigenomic patterns, we analyzed ChIP-seq data from Roadmap and ENCODE projects, including active (H3K4me3, H3K9ac, and H2A.Z) and repressive (H3K27me3) promoter modifications, and marks associated with enhancers (H3K4me1, H3K27ac, DNase I hypersensitivity sites). We used the width of called ChIP-seq peaks for promoter features and counted the interacting number of promoters and enhancers within pre-defined topologically-associated domains (TADs) for enhancer features. As each histone modification is characterized in multiple cell types, we refer to the combination of an epigenomic modification and a cell type as one epigenomic feature.

Figure 1A shows the correlation among epigenomic features, and the correlation of epigenomic features and ExAC pLI score. As expected, active promoter or enhancer marks are highly correlated with each other and with ExAC pLI score, and they are anti-correlated with repressor marks in general. The repressor marks from stem cells or fetal tissues have positive correlations with active marks and ExAC pLI scores, suggesting many genes with bivalent marks in stem cells are likely haploinsufficient.

**Figure 1.**
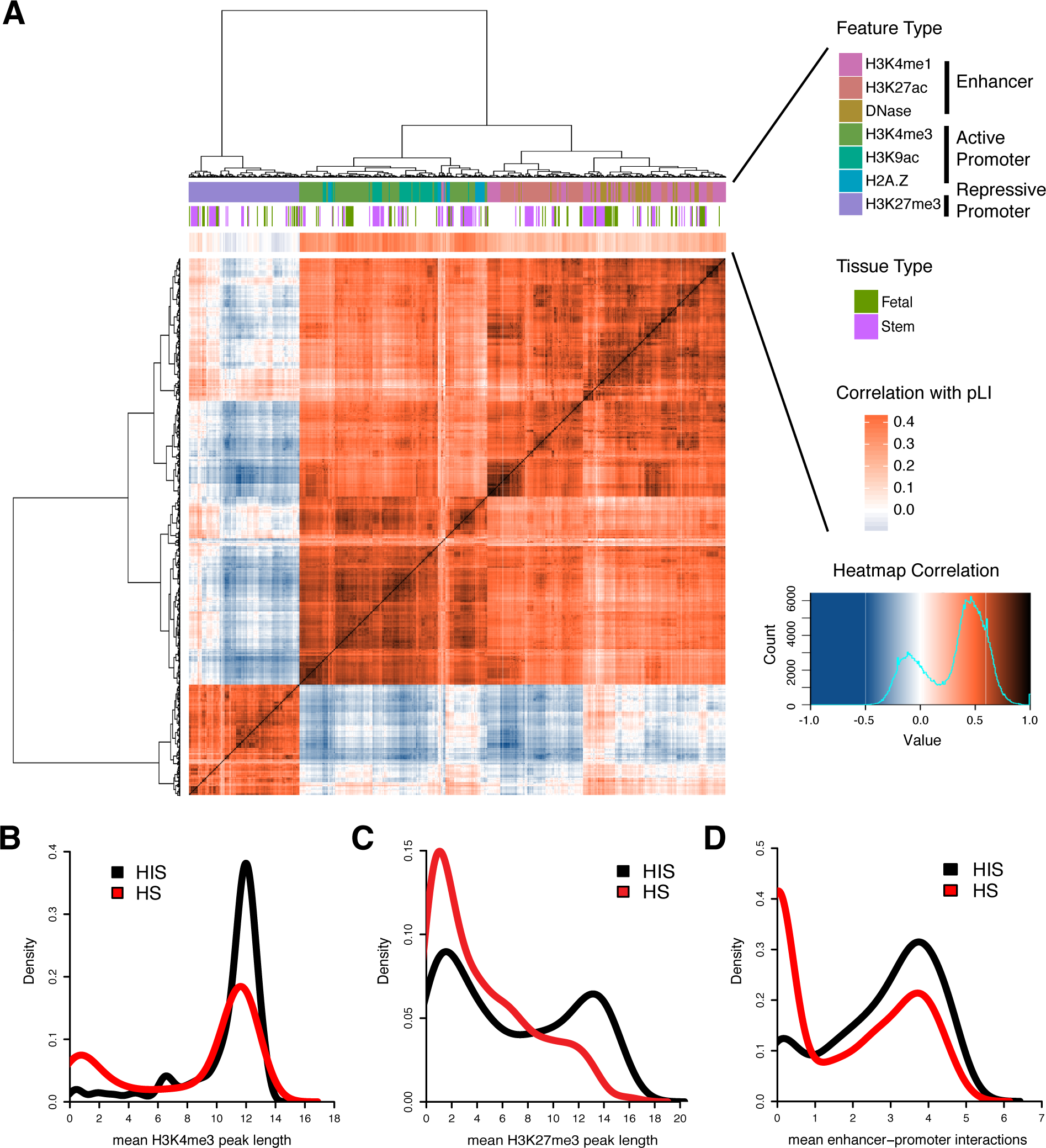
Epigenomic profiles are associated with gene haploinsufficiency. (A) Heatmap showing Spearman correlation between epigenomic features. Three groups of epigenomic features are included: active promoter, repressive promoter and enhancer features. Epigenomic features inside each group strongly correlate with each other. Different feature types, including various histone modifications, histone variant, and DNase I hypersensitivity sites, are color-coded. Above the heatmap, a bar denoting Spearman correction between epigenomic features and pLI shows many epigenomic features relate to HIS with varying degree. Data from stem cells or fetal tissues are also marked by color lines. (B-C) Known HIS and HS genes have different distributions of peak length of promoter features (B, H3K4me3; C, H3K27me3). For each gene, peak length was averaged across tissues. (D) HIS and HS genes have different distributions of number of interacting enhancers inferred by Epitensor. For each gene, the number of interacting enhancers was averaged across tissues.

To further investigate the association of haploinsufficiency and patterns of epigenomic modifications, we compiled a list of 287 known HIS genes (Supplementary Table 1) involved in a wide range of human diseases (Supplementary Table 2) from a recent study ^8,18^ and human-curated ClinGen dosage sensitivity map. We also collected a list of 717 HS genes, of which one copy of each gene had been deleted in two or more subjects based on a CNV study in 2,026 healthy individuals ^19^. For promoter features, HIS and HS genes clearly have distinct distributions of peak length (Figure 1B-D). HIS genes on average have wider peaks of both the active marker H3K4me3 (Figure 1B) and the repressive marker H3K27me3 (Figure 1C), suggesting the difference between HIS and HS genes is not only on the level of expression but also on distinct mechanisms of regulation. Furthermore, other epigenomic modifications associated with active promoters, including H2A.Z and H3K9ac, also display wider peaks upstream of HIS genes (Supplementary Figure 1 A and B). In addition, HIS and HS genes also differ in the number of interacting enhancers. We adopted a recently published method *EpiTensor* ^20^, which decomposes a 3D tensor representation of histone modifications, DNase-Seq, and RNA-Seq data to find associations between distant genomic regions. When restricted to pre-defined topologically-associated domains (TADs), associated regions identified by EpiTensor correspond well to enhancer-promoter interactions found by Hi-C. EpiTensor revealed that HIS genes have a median of 9 interacting enhancers, while HS genes have a median of 0 (p < 10^-4^, permutation test, Supplementary Figure 1C). When averaged across tissues, HIS genes shift towards a larger number of mean interacting enhancers, as compared to HS genes (Figure 1D), supporting the notion that HIS genes have more regulatory complexity.

Among these 287 known HIS genes, 129 genes (45%) have pLI smaller than 0.9 or missing value. Some of these genes are well-known disease risk genes under dominant genetic models, such as *TGFB1*^21^, *RUNX1*^22^, *SoX2*^23^, *SUMO1*^24^, *NKX2-5*^25^, *EYA4*^26^, *CAV1*^27^, *PAX2*^28^, *GATA6*^29^, *ZIC2*^30^, and *WT1*^31^. These known HIS genes with pLI < 0.9 have significantly smaller number of expected loss of function variants^11^ than an average gene (Supplementary Figure 2A), and intermediate selection coefficient (S_het_) ^12^ (Supplementary Figure 2B), pointing to two particular areas (genes that are either short or under intermediate negative selection) in which HIS prediction can be improved.

**Figure 2.**
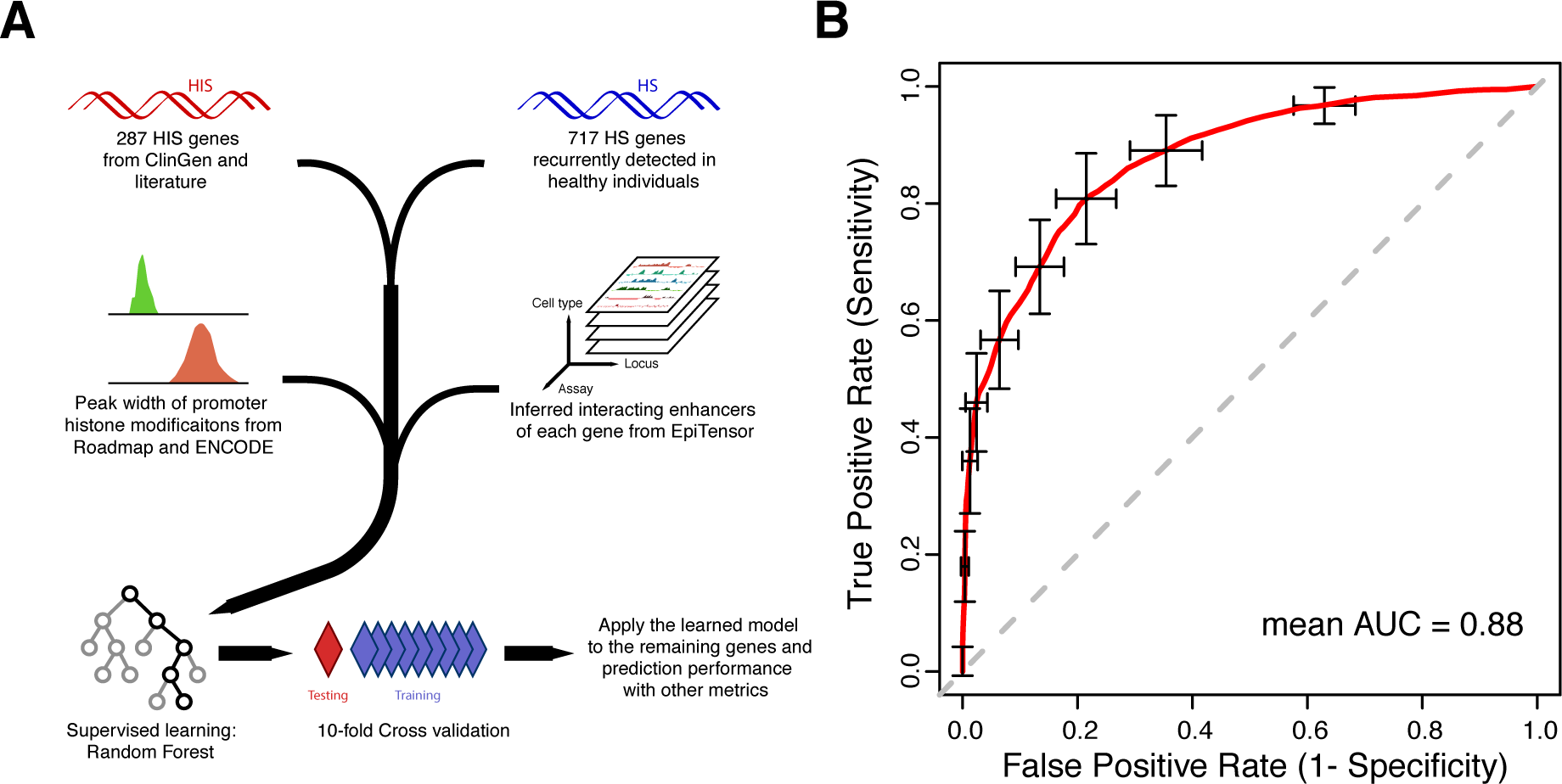
A Random Forest model to predict haploinsufficiency. (A) A flowchart of the method. (B) ROC curve from 10-fold cross-validation. The red curve is the average of 100 randomized cross-validation runs, with error bar showing standard deviation. The mean and median AUC of the 100 runs are 0.88 and 0.89, respectively. (Supplementary Table 4). Similarly to the ones with low pLI values in the positive training set, these genes have lower background mutation rate (which is primarily determined by transcript size) than the ones with large pLI values (Supplementary Figure 3D), and are generally under less severe selection measured by S_het_ ^12^ (Supplementary Figure 3E).

### Predicting haploinsufficiency with epigenomic features

To leverage the strong association between epigenomic patterns and gene haploinsufficiency, we developed a computational method to predict haploinsufficiency using Random Forest (Figure 2A) and other supervised learning models (Supplementary Figure 3 A and B). The input features included peak length of four promoter marks (H3K4me3, H3K9ac, H2A.Z and H3K27me3) and the number of EpiTensor-inferred interacting enhancers in various tissues. Performance evaluation by 10-fold cross validation and AUC (Area Under Curve) in ROC (Receiver Operating Characteristic) curves showed that all of these methods achieved high AUC values of 0.86~0.88 (Figure 2B and Supplementary Figure 3 A and B). As Random Forest performs the best, results from Random Forest are chosen as final metrics measuring the probability of being haploinsufficient, termed “Episcore” (Supplementary Table 3). Despite completely different input data are used, Episcore and ExAC pLI score displayed overall concordance. The distribution of pLI is generally bi-modal, with modes at 1 and 0 ^11^. The genes with Episocre >0.6 are much more likely to have pLI values close to 1 than genes with Episocre < 0.4, and the opposite trend at pLI close to 0 (Supplementary Figure 3C). Among 3463 genes with Episcore > 0.6, 1518 have pLI scores < 0.5. Some of these genes have been implicated in human diseases under a dominant model, such as *HEY2*^32^, *ASF1A*^33^ and *HAND2*^34^

### Episcore provides better prioritization of *de novo* LGD variants in developmental disorders

A major goal of predicting haploinsufficiency is to facilitate prioritization of variants identified in genetic studies of developmental disorders. We compared Episcore with pLI scores from ExAC ^11^, S_het_ values (selection coefficient of heterozygous LGD variants) ^12^, and ranks of mouse heart expression level ^35^, using *de novo* LGD variants identified in a recently published whole exome sequencing study DDD (Deciphering Developmental Disorders consortium) of 1,365 trio families with congenital heart disease (CHD) ^36^. LGD variants include frameshift, nonsense and canonical splice site mutations. We only included genes with all 4 metrics for comparison, although we note Episcore (19,430 genes) made predictions for more genes than pLI (18,225 genes), S_het_ (17,200 genes) and ranks of mouse heart expression level (17,624 genes, due to loss in orthologue matching). Different predictions are compared by the enrichment rate of variants. For the same number of top-ranked genes by each metric, we calculated the number of LGD variants located in these genes and estimated the number of LGD variants based on background mutation rate ^37^. Across a wide range of top-ranked genes, Episcore showed larger enrichment than ExAC pLI, S_het_, or heart expression level (Figure 3A and Supplementary Figure 4A). We also applied the same approach to *de novo* synonymous variants identified in the CHD dataset and observed no enrichment (Supplementary Figure 4B). Additionally, we compared these predictions by precision-recall-like curve (PR-like) based on enrichment. Since the total number of positive variants (true disease-causing variants) is unknown, we used estimated number of “true positives” instead of “true positive rate (recall)” in this comparison. For top-ranked genes from each method, the number of true positives were estimated by subtracting expected number of LGD variants based on background mutation rate from the observed in these genes. We measured precision by dividing the estimated number of true positives by the total number of observed LGD variants in these genes. Across a wide range of precision, Episcore consistently showed superior recall compared to pLI, S_het_ and heart expression level (Figure 3B) and to earlier methods based on combination of genetic and protein interaction network data ^8,9^ (Supplementary Figure 4C and D). The performance advantage over other HIS-related score does not change after excluding the genes used in training (Supplementary Figure 4E and F).

**Figure 3.**
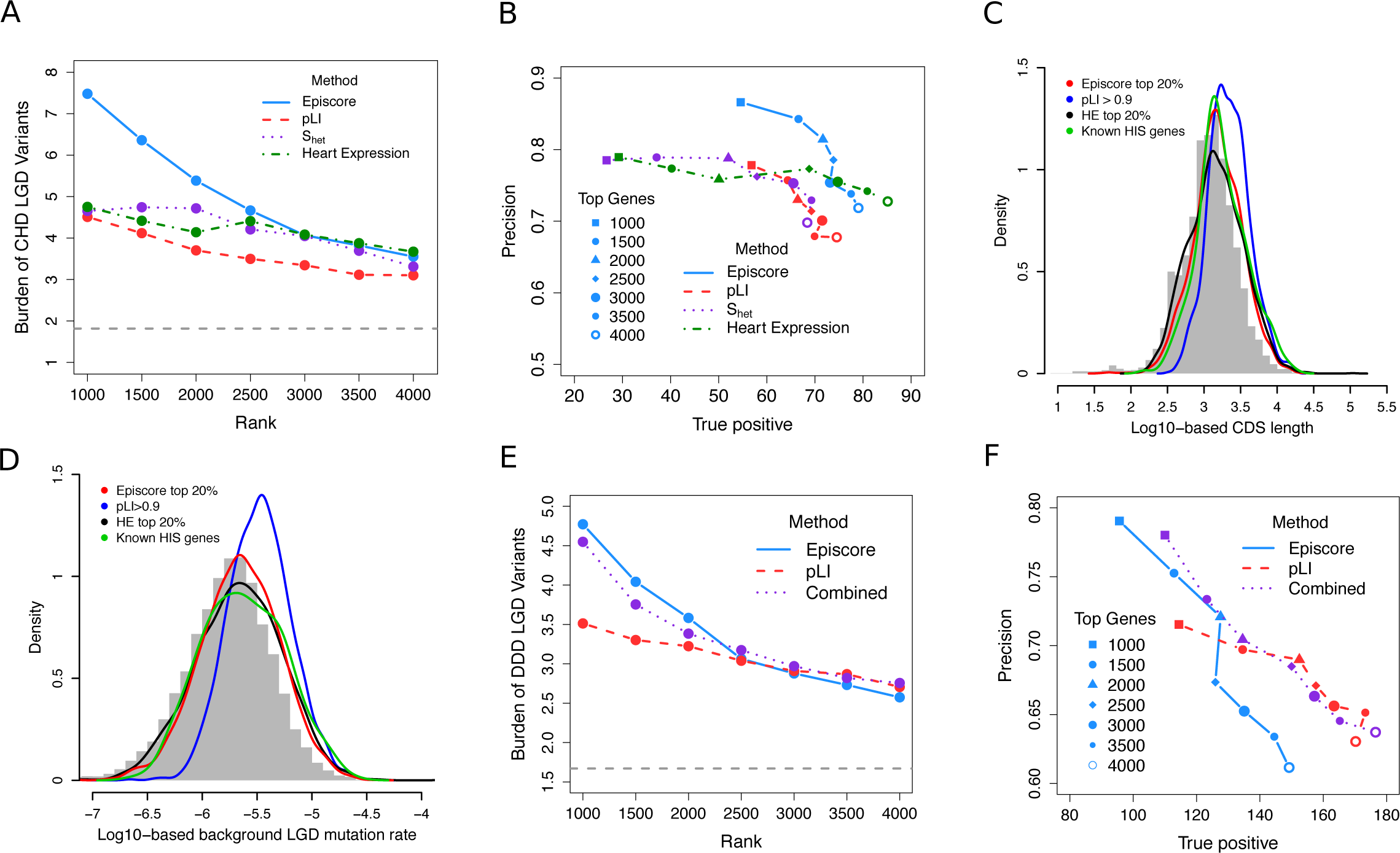
Assessment of the performance of Episcore in variant prioritization using *de novo* mutation data. (A-B) Comparison of Episcore, pLI, S_het_ and heart expression level (HE) in variant prioritization using CHD exome sequencing data. In (A), burden refers to the ratio between the number of *de novo* LGD variants observed in top genes ranked by each metric and the number of expected *de novo* LGD variants due to background mutation. Episcore has higher enrichment in top 1000-2500 genes and similar enrichment afterwards. The grey dash line indicates the burden of *de novo* LGD variants in all genes. (B) Precision-recall-like curves. True positive is the difference between the observed and expected *de novo* LGD variants. Precision is calculated by dividing the number of true positives by the number of observed *de novo* LGD variants. The blue curve for Episcore shifts upright than pLI and S_het_, showing Episcore has better recall with precision and vice versa. (C-D) Episcore has less bias towards genes with longer CDS length (C) or larger background mutation rate (D) than pLI. Grey histogram in the background represents CDS length or mutation rate of all genes in the genome. The blue curve for pLI shifts right, while the curves for Episcore and HE are similar to the distribution of all genes and known HIS genes. (E-F) A combination of Episcore and pLI, the meta-score, has better performance in variant prioritization when benchmarked using DDD exome sequencing data. Meta-score is the output from a logistic regression model, using Episcore and pLI as input. Enrichment, true positive and precision were calculated similarly to (A-B).

**Figure 4.**
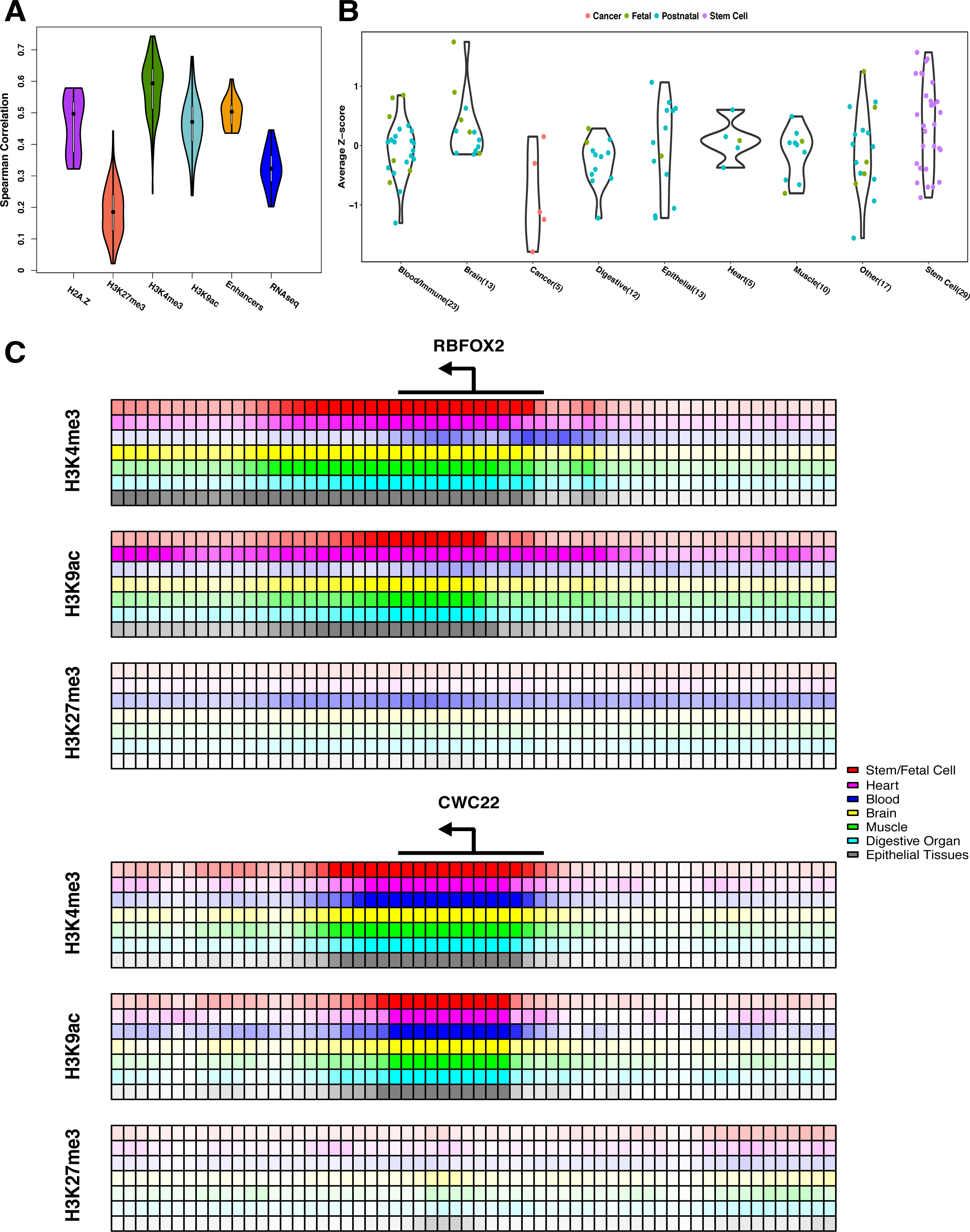
Contribution of epigenomic features to Episcore prediction. (A) Spearman correlation between epigenomic feature and Episcore. Features used in the Random Forest model, including H2A.Z, H3K27me3, H3K4me3, H3K9ac and the number of interacting enhancers, all have positive correlation with Episcore. Spearman correlation coefficients between gene expression level, measured in RPKM (reads per kilobase per million reads), and Episcore were also plotted for comparison. (B) The importance of each tissue in generating Episcore is measured by average Z-score, which is converted from Spearman correlation coefficients between epigenomic feature and Episcore. Each dot represents one cell line or tissue type indicated by colors. Stem cells and neural and fetal tissues are the most important tissue and cell types in Episcore prediction. (C) The epigenomic profile of an example HIS gene, *RBFOX2*, and a house-keeping gene, *CWC22*. Each small box represents 100bp region around TSS and the shade of the color reflects averaged fold change of reads between ChIP-seq library and control samples. *RBFOX2* has a broad expansion of epigenomic marks while *CWC22* is not, and *RBFOX2* shows more tissue-specific regulation but *CWC22* has narrow peaks in active marks across all the tissues.

We obtained a second CHD WES cohort of 2,645 parent-offspring trios from the Pediatric Cardiac Genomics Consortium (PCGC) ^38^ to emulate a replication design. We used the larger data (PCGC CHD) as discovery and the DDD data as replication. We found that the genes with a single LGD variant in PCGC data and “replicated” with at least one LGD variant in the DDD data have much higher Episcore, than the genes with a singleton LGD in PCGC data or genes with LGD variants in controls (unaffected siblings in Simons Simplex Collection autism study ^39^)(Supplementary Figure 5).

### Episcore provides complementary information to mutation intolerance metrics

Haploinsufficiency predicted by mutation intolerance in a general population (such as ExAC pLI metric) is intrinsically biased towards genes with longer CDS (coding sequence) lengths or higher background mutation rates. Figure 3C and D show the distribution of genes with pLI scores > 0.9 shifts towards longer CDS length or higher background mutation rate, as compared to the distribution of known HIS disease risk genes, while top 20% genes ranked by Episcore have similar distribution to known HIS disease risk genes or genes with expression level ranked in top 20% in developing heart ^35^.

Since Episcore and pLI use distinct types of input data, a combination of these two scores might achieve better performance. We obtained de novo mutation data of 4,293 trio families affected by developmental disorders, mostly with intellectual disabilities (DDD ID), from a recent study^6^. Genes with *de novo* LGD mutations in DDD ID cases are notably under more severe selection than the ones in CHD (Supplementary Figure 4G). We used a logistic regression to integrate Episcore and pLI in this data set. Specifically, we used a total of 45 genes with *de novo* LGD variants in 3 or more probands as positives, and randomly sampled 45 genes from genes with no observed *de novo* LGD variant as negatives to estimate coefficients in the logistic model. Both Episcore and pLI have significant coefficients (P < 10^-3^), supporting these two methods convey complementary information. We found that the resulting meta-score achieved overall better precision and true positives than Episcore or pLI alone (Figure 3 E and F), while maintaining similar enrichment burden as good as any method alone in a broad range of gene ranks.

### Brain tissues, fetal tissues, and stem cells highly associate with the predicted haploinsufficiency

To evaluate the association of each epigenomic feature to haploinsufficiency, we calculated Spearman correlation coefficients between each feature and Episcore. These correlation coefficients were analyzed in two ways. We first grouped them based on the molecular entities they represent, such that the same epigenomic modification from different tissues would be in one group. Each of the 5 resulting categories has distinct distributions of Spearman correlation coefficients, suggesting different contributions to Episcore (Figure 4A). Except for the repressive mark H3K27me3, most of them have larger correlation coefficients than gene expression values, suggesting these features and the model do not merely reflect expression abundance but also epigenomic regulation specific to HIS genes. Measured by mean decrease of Gini index, these groups of features have similar trend in contribution to Episcore prediction (Supplementary Figure S6).

We then grouped correlation coefficients based on tissue and cell types, converted correlation coefficient of each epigenomic modification to a Z-score using the mean and standard deviation across the tissue or cell type, and finally averaged the Z-score of all epigenomic modification for each tissue or cell type. The averaged Z-score represents the importance of this tissue or cell type to haploinsufficiency prediction. In general, stem cells and neural tissues have large average Z-scores (Figure 4B). Interestingly, for tissues in the same category, fetal tissues usually have larger average Z-scores than postnatal tissues.

Finally, to illustrate the contribution of different tissues to HIS, we examined in detail the histone modifications around TSS of several known HIS genes. Figure 4C show *RBFOX2* and *CWC22. RBFOX2* is a CHD risk gene recently discovered through *de novo* LGD variants ^5^, and it has expansive H3K4me3 and H3K9ac peaks in stem/fetal cells and heart and brain tissues, but not in blood cells. Consistently, it has a reverse pattern in H3K27me3, extensive in blood cells but limited in other tissues. On the contrary, *CWC22*, a known house-keeping gene, shows consistent but narrow peaks of active marks across tissues.

## Discussion

In this study we showed there is a strong correlation between epigenomics patterns and gene haploinsufficiency, and developed a computational method (Episcore) to predict HIS using epigenomic features. Episcore had superior yet complementary performance in prioritization of *de novo* LGD variants in congenital heart disease and neurodevelopmental disorders, compared to mutation intolerance metrics such as ExAC pLI ^11^.

Existing HIS prediction methods based on intolerance of mutations have limited statistical power in genes with small transcript size or under less severe negative selection. Network-based methods^8^ are often biased towards well-studied genes ^9^ and pathways. Epigenomic data have several advantages to address these issues: (a) they are orthogonal to genetic mutations, and therefore provide additional information that could improve power; (b) they are much less biased by transcript size, and will be most helpful to predict HIS of genes with short transcripts; (c) the bias with selection coefficient is a reflection of the training data, which empirically is much smaller than mutation intolerance metrics; (d) the ability to generate large amount of data without bias towards well-studied genes. These advantages contribute to the superior performance of Episcore in prioritizing *de novo* LGD variants from exome sequencing studies.

There are likely a variety of mechanisms underlining the correlation of epigenomics patterns and haploinsufficiency. First, broad H3K4me3 peaks contributed most to Episcore prediction of HIS. Broad H3K4me3 peaks are associated with reduced transcriptional noise at cell population and single cell levels ^15^, which is likely required to maintain precise expression levels of HIS genes in specific cell types and developmental stages. Second, a previous study found regulatory complexity is required to achieve cell-type specific expression patterns of the lineage-defining genes in hematopoietic differentiation ^40^. Consistently, we found the number of enhancers interacting with the promotor of a gene is highly correlated with predicted HIS score. Third, many HIS genes are regulators that define cell lineages during differentiation. Bivalent chromatin domains in embryonic stem cells, in which both active marker H3K4me3 and repressor marker H3K27me3 are present, are generally associated with lineage control genes ^41^. We observed that H3K27me3 are positively correlated with H3K4me3 in stem cells, and both are correlated with mutation intolerance (Figure 1A and 1C) and Episcore (Figure 4A). Finally, we found epigenomic features from stem cells and fetal tissues contribute most to prediction, highlighting the importance of developmental role in determining gene haploinsufficiency.

Our data suggests Episcore is generally better for prioritizing genes with a broader range of selection coefficient or genes with smaller transcript size, whereas pLI performs better for genes under most severe negative selection. Episcore is currently limited by availability and resolution of epigenomic data, especially cell-type specific data from complex tissues or organs such as the brain, and data at various developmental stages. Complex developmental disorders, such as autism, involve a large number of cell types during a broad range of developmental stages. It is critical to generate and integrate more fine-grained epigenomic data from cells of specific types at specific time points in order to improve genetic discoveries in studies of such diseases. We expect such data sets will become available in near future from ongoing projects ^42–44^, and will enable us to improve prediction of HIS and facilitate novel discoveries in genetic studies.

## Methods

### Collection and Preprocessing of Training Genes

In this study, we used Ensembl release 75 for gene annotation and TSS (transcription start site) locations. All genomic coordinates are based on hg19 human genome assembly. Any non-hg19 coordinates were lifted over to hg19 using UCSC LiftOver tool (https://qenome.ucsc.edu/cqi-bin/hqLiftOver). Conversion of gene symbols to Ensembl IDs were based on annotation tables downloaded from Ensembl BioMart.

Positive training set data (curated haploinsufficient genes) were collected from these two sources: (1) haploinsufficent training genes used in previous studies ^8,18^ and (2) genes with haploinsufficient score of 3 in ClinGen Dosage Sensitivity Map (http://www.ncbi.nlm.nih.gov/proiects/dbvar/clingen/). For the negative training set (curated haplosufficient genes), we used genes deleted in two or more healthy people, based on CNVs detected in 2,026 normal individuals ^19^. Only genes with half or more of its length covered by any deletion were considered “deleted” in an individual.

The raw training set may have some false positives and false negatives, as it contained results from automated literature mining that is known to give noisy output. To optimize the performance, we did the following pruning of the raw training set: (1) we only kept protein-coding genes in autosomes, as non-protein-coding genes or genes on sex chromosomes may be under different mechanism of epigenomic regulation; (2) from the positive training set, we removed genes with sufficient contradictory evidence (ExAC pLI ≤ 0.1 and expected loss-of-function variants > 10 ^11^); and (3) from the negative training set, we removed genes with sufficient contradictory evidence (pLI ≥ 0.9 and expected loss-of-function variants > 10). After pruning, the positive training set has 287 genes and the negative training set has 717 genes. The full list of training genes is available in Supplementary Table 1.

### Preprocessing of Epigenomic Feature Data

The uniformly processed peak calling results of Roadmap and ENCODE projects were downloaded from http://egg2.wustl.edu/roadmap/web_portal/processed_data.html. For promoter features (H2A.Z, H3K27me3, H3K4me3, and H3K9ac), “GappedPeaks” were used to allow for broad domains of ChIP-seq signal. The assignment of a GapppedPeak to a gene follows these steps in order: (1) for each gene, only TSS of Ensembl canonical transcripts were used. (2) assigned a GappedPeak to a TSS if the GappedPeak overlaps with the upstream 5kb to downstream 1kb region around the TSS. This definition of basal cis-regulatory region around promoter follows GREAT tool ^45^. Assigning one GappedPeak to multiple TSS was allowed. (3) For TSS having more than 1 GappedPeak assigned, kept the closest one. (4) For genes with multiple TSS and hence multiple assigned GappedPeaks, kept the longest GappedPeak. After these four steps, if one gene had been associated with a GappedPeak, then we used the width of the peak as an epigenomic feature in the following machine learning models. If a gene had no associated GappedPeak, then the peak width is 0.

To calculate the number of interacting enhancers of a gene, we used two approaches. In a naïve approach, we counted peaks of ChIP-seq signals that are associated with enhancers. The ChIP-seq signals we used include H3K4me1, H3K27ac and DNase I hypersensitivity site, and each ChIP signal was counted and recorded separately. We used “NarrowPeak” instead of “GappedPeak” in the counting to better estimate the number of interacting enhancers, as enhancer regions are not long and GappedPeak has the risk of merging nearby ChIP-seq signals. For each gene, we counted peaks in (a) the surrounding TAD (Topologically Associated Domain), based on TADs reported in ^46^; or (b) +/-20kb of each TSS (Only TSSs of Ensembl canonical transcripts were used. For genes with multiple TSS and thus several numbers of interacting enhancers, we kept the largest one). In a more advanced approach, we adapted EpiTensor ^20^ to infer gene-enhancer relationship. We made a few changes when using EpiTensor: (a) we used normalized coverage of ChIP-seq signal instead of raw coverage in Zhu et al. 2016 ^20^; (b) we used the coverage of H3K27ac, H3K27me3, H3K36me3, H3K4me1, H3K4me3, H3K9me3, DNase I and RNA-seq as input for EpiTensor to balance between more input data types and more cell types included, as not every cell type has all these histone modifications characterized. The number of data types included are fewer than the ones used in Zhu et al. 2016 ^20^, but it could still achieve desirable performance (personal communications); (c) we used enhancer annotation from 15-state chromHMM (http://egg2.wustl.edu/roadmap/web_portal/chr_statelearning.html#core_15state), while the original EpiTensor paper ^20^ used results of an earlier version. Based on the output of EpiTensor, which predicts enhancer-promoter pairs, we counted the number of interacting enhancers for each gene in various tissues.

Finally, the results of peak width and number of interacting enhancers were consolidated into a matrix, with each row being a gene and each column representing a combination of a tissue and a data type, e.g. “H3K4me3 peak width in fetal heart”. One combination of a tissue and a data type was referred to as one epigenomic feature. This matrix was used as input for machine learning models described in the following section.

### Machine learning approaches to predict haploinsufficiency

We applied several machine learning approaches, including Random Forest, Support Vector Machine (SVM) and SVM with LASSO feature selection. Random Forest was implemented using R package “randomForest”. SVM was implemented using R package “e1071”. LASSO was implemented using R package “glmnet”, with alpha value equal to 1. For each machine learning method, we assessed the performance based on 100 runs of 10-fold cross-validation. In each run, 10% of the training genes were randomly selected and left out to form a test set for validation. The remaining data were used to train the model, after which the test set was used to calculate model sensitivity and specificity. We used R package “ROCR” to make an ROC curve based on the 100 runs and calculated AUC values.

Finally, we used all training genes used to train the model, and then estimate the probabilities of being positive (i.e. probabilities of being HIS) for all genes. The whole process was repeated 30 times and we took the arithmetic mean of the 30 sets of probabilities as the final results.

### Comparing Episcore and other metrics in variant prioritization

We used two approaches to compare Episcore and other metrics in variant prioritization, based on “enrichment of *de novo* LGD variants”, estimated “number of true-positives” and “precision”. The formula to calculate these three statistics are as follows.

For any gene *i*, the number of expected *de novo* LGD variants in each gene, ***E**_i_*, was calculated as:

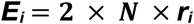

where ***N*** is the number of cases in the sequencing cohort and *r_i_* is gene-specific LGD mutation rate. LGD variants include nonsense, frameshift and canonical splice site mutations. The background mutation rate per gene of each mutation type was obtained from Samocha et al. 2014 ^37^. For each gene, ***r**_i_* is the sum of background mutation rate of nonsense, frameshift and canonical splice site mutations.

For a set of genes, the enrichment of *de novo* LGD variants, ***D***, was calculated as:

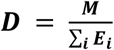

where ***M*** is the total number of observed *de novo* LGD variants in this gene set. In this study, we used results from two whole exome sequencing studies on congenital heart disease ^5,36^ and another whole exome sequencing study on various developmental disorders ^6^.

For any gene set, the number of true positives, ***TP***, was calculated as:

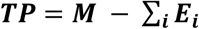

For any gene set, the precision (positive predictive value), ***PPV***, was calculated as:

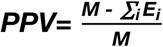

For each metric (Episcore, pLI, etc.), a series of top-ranked genes were selected, such as top 500 genes, top 2000 genes, etc. In the first approach, enrichment of *de novo* LGD variants, **D**, was calculated for any set of top-ranked genes, and then enrichment values were plotted and compared, as shown in Figure 3A. In the second approach, the number of true positives, ***TP***, and the precision (true discovery rate), ***PPV***, were calculated for any set of top-ranked genes. ***TP*** and ***PPV*** were plotted and compared, as shown in Figure 3B. If the number of all true positives (**N**) in a study is known, we can calculate recall as *R* = *TP/N*. Although *N* is generally unknown, it is a constant; therefore, *TP* is proportional to R. In this study, we use TP as a proxy of recall.

To examine the utility of Episcore in prioritizing genes with only one LGD mutation, we utilized two independent Congenital Heart Disease (CHD) cohorts: DDD (Deciphering Developmental Disorders consortium) CHD ^36^ and PCGC (Pediatric Cardiac Genomics Consortium) CHD ^38^. Both these two study included trios from an earlier CHD study ^35^ to increase detection power. To avoid duplication, we removed these earlier trios from DDD CHD data.

### Epigenomic features critical in the prediction

We calculated a Spearman correlation coefficient between each epigenomic feature and Episcore. One epigenomic feature here corresponds to a data type (like H3K4me3 peak width) in certain tissue/cell type (e.g. fetal heart). To examine which data types are more important, we plotted these Spearman correlation coefficients by data type, e.g. correlation coefficients from H3K4me3 peak width were plotted in one section. To examine what tissue/cell types are more important, we calculated averaged z-score for each tissue/cell type. The average z-score is calculated following these two steps: (1) we converted every Spearman correlation coefficient to a Z-score using mean and standard deviation specific to each data type and (2) for each tissue/cell type, we averaged the Z-scores from various data types.

## Acknowledgements

We thank Dr. Zhao Chen for advices on setting up and running EpiTensor. The work was partly supported by NIH grant R01GM120609 (S.C. and Y.S.).

## Author Contributions

All authors contributed to data analysis, interpretation, and manuscript writing. X.H. and S.C. curated and processed epigenomics data sets. Y.S. conceived and designed the study.

## References

1. De Rubeis, S. et al. Synaptic, transcriptional and chromatin genes disrupted in autism. Nature 515, 209–15 (2014).

2. Iossifov, I. et al. The contribution of de novo coding mutations to autism spectrum disorder. Nature 515, 216–21 (2014).

3. Hamdan, F.F. et al. De novo mutations in moderate or severe intellectual disability. PLoS Genet 10, e1004772 (2014).

4. Deciphering Developmental Disorders, S. Large-scale discovery of novel genetic causes of developmental disorders. Nature 519, 223–8 (2015).

5. Homsy, J. et al. De novo mutations in congenital heart disease with neurodevelopmental and other congenital anomalies. Science (New York, N.Y.) 350, 1262–6 (2015).

6. McRae, J.F. et al. Prevalence, phenotype and architecture of developmental disorders caused by de novo mutation. bioRxiv (2016).

7. He, X. et al. Integrated model of de novo and inherited genetic variants yields greater power to identify risk genes. PLoS Genet 9, e1003671 (2013).

8. Huang, N., Lee, I., Marcotte, E.M. & Hurles, M.E. Characterising and predicting haploinsufficiency in the human genome. PLoS Genet 6, e1001154 (2010).

9. Steinberg, J., Honti, F., Meader, S. & Webber, C. Haploinsufficiency predictions without study bias. Nucleic Acids Res 43, e101 (2015).

10. Petrovski, S., Wang, Q., Heinzen, E.L., Allen, A.S. & Goldstein, D.B. Genic intolerance to functional variation and the interpretation of personal genomes. PLoS Genet 9, e1003709 (2013).

11. Lek, M. et al. Analysis of protein-coding genetic variation in 60,706 humans. Nature 536, 285–91 (2016).

12. Cassa, C.A. et al. Estimating the selective effects of heterozygous protein-truncating variants from human exome data. Nat Genet 49, 806–810 (2017).

13. Chen, K. et al. Broad H3K4me3 is associated with increased transcription elongation and enhancer activity at tumor-suppressor genes. Nat Genet (2015).

14. Davoli, T. et al. Cumulative haploinsufficiency and triplosensitivity drive aneuploidy patterns and shape the cancer genome. Cell 155, 948–62 (2013).

15. Benayoun, B.A. et al. H3K4me3 breadth is linked to cell identity and transcriptional consistency. Cell 158, 673–88 (2014).

16. Roadmap Epigenomics, C. et al. Integrative analysis of 111 reference human epigenomes. Nature 518, 317–30 (2015).

17. ENCODE Project Consortium. An integrated encyclopedia of DNA elements in the human genome. Nature 489, 57–74 (2012).

18. Dang, V.T., Kassahn, K.S., Marcos, A.E. & Ragan, M.A. Identification of human haploinsufficient genes and their genomic proximity to segmental duplications. Eur J Hum Genet 16, 1350–7 (2008).

19. Shaikh, T.H. et al. High-resolution mapping and analysis of copy number variations in the human genome: a data resource for clinical and research applications. Genome Res 19, 1682–90 (2009).

20. Zhu, Y. et al. Constructing 3D interaction maps from 1D epigenomes. Nat Commun 7, 10812 (2016).

21. Kinoshita, A. et al. Domain-specific mutations in TGFB1 result in Camurati-Engelmann disease. Nat Genet 26, 19–20 (2000).

22. Taketani, T. et al. Mutation of the AML1/RUNX1 gene in a transient myeloproliferative disorder patient with Down syndrome. Leukemia 16, 1866–7 (2002).

23. Fantes, J. et al. Mutations in SOX2 cause anophthalmia. Nat Genet 33, 461–3 (2003).

24. Alkuraya, F.S. et al. SUMO1 haploinsufficiency leads to cleft lip and palate. Science 313, 1751 (2006).

25. Benson, D.W. et al. Mutations in the cardiac transcription factor NKX2.5 affect diverse cardiac developmental pathways. J Clin Invest 104, 1567–73 (1999).

26. Wayne, S. et al. Mutations in the transcriptional activator EYA4 cause late-onset deafness at the DFNA10 locus. Hum Mol Genet 10, 195–200 (2001).

27. Cao, H., Alston, L., Ruschman, J. & Hegele, R.A. Heterozygous CAV1 frameshift mutations (MIM 601047) in patients with atypical partial lipodystrophy and hypertriglyceridemia. Lipids Health Dis 7, 3 (2008).

28. Sanyanusin, P. et al. Mutation of the PAX2 gene in a family with optic nerve colobomas, renal anomalies and vesicoureteral reflux. Nat Genet 9, 358–64 (1995).

29. Kodo, K. et al. GATA6 mutations cause human cardiac outflow tract defects by disrupting semaphorin-plexin signaling. Proc Natl Acad Sci U S A 106, 13933–8 (2009).

30. Brown, S.A. et al. Holoprosencephaly due to mutations in ZIC2, a homologue of Drosophila odd-paired. Nat Genet 20, 180–3 (1998).

31. Hastie, N.D. Dominant negative mutations in the Wilms tumour (WT1) gene cause Denys-Drash syndrome--proof that a tumour-suppressor gene plays a crucial role in normal genitourinary development. Hum Mol Genet 1, 293–5 (1992).

32. Reamon-Buettner, S.M. & Borlak, J. HEY2 mutations in malformed hearts. Hum Mutat 27, 118 (2006).

33. Giannakou, A. et al. Copy number variants in Ebstein anomaly. PLoS One 12, e0188168 (2017).

34. Sun, Y.M. et al. A HAND2 Loss-of-Function Mutation Causes Familial Ventricular Septal Defect and Pulmonary Stenosis. G3 (Bethesda) 6, 987–92 (2016).

35. Zaidi, S. et al. De novo mutations in histone-modifying genes in congenital heart disease. Nature 498, 220–3 (2013).

36. Sifrim, A. et al. Distinct genetic architectures for syndromic and nonsyndromic congenital heart defects identified by exome sequencing. Nat Genet 48, 1060–5 (2016).

37. Samocha, K.E. et al. A framework for the interpretation of de novo mutation in human disease. Nat Genet 46, 944–50 (2014).

38. Jin, S.C. et al. Contribution of rare inherited and de novo variants in 2,871 congenital heart disease probands. Nat Genet (2017).

39. Krumm, N. et al. Excess of rare, inherited truncating mutations in autism. Nat Genet 47, 582–8 (2015).

40. Gonzalez, A.J., Setty, M. & Leslie, C.S. Early enhancer establishment and regulatory locus complexity shape transcriptional programs in hematopoietic differentiation. Nat Genet (2015).

41. Vastenhouw, N.L. & Schier, A.F. Bivalent histone modifications in early embryogenesis. Curr Opin Cell Biol 24, 374–86 (2012).

42. Stunnenberg, H.G., International Human Epigenome, C. & Hirst, M. The International Human Epigenome Consortium: A Blueprint for Scientific Collaboration and Discovery. Cell 167, 1145–1149 (2016).

43. Psych, E.C. et al. The PsychENCODE project. Nat Neurosci 18, 1707–12 (2015).

44. Dekker, J. et al. The 4D Nucleome Project. bioRxiv (2017).

45. McLean, C.Y. et al. GREAT improves functional interpretation of cis-regulatory regions. Nat Biotechnol 28, 495–501 (2010).

46. Dixon, J.R. et al. Topological domains in mammalian genomes identified by analysis of chromatin interactions. Nature 485, 376–80 (2012).

